# Mesenchymal ETV Transcription Factors Regulate Cochlear Length

**DOI:** 10.1101/2020.05.01.072454

**Authors:** Michael Ebeid, Sung-Ho Huh

**Affiliations:** Department of Neurological Sciences, University of Nebraska Medical Center, Omaha, NE, 68198, USA; Holland Regenerative Medicine Program, University of Nebraska Medical Center, Omaha, NE, 68198, USA

**Keywords:** cochlear development, FGF, ETV transcription factors, gene expression

## Abstract

Mammalian cochlear development encompasses a series of morphological and molecular events that results in the formation of a highly intricate structure responsible for hearing. One remarkable event occurs during development is the cochlear lengthening that starts with cochlear outgrowth around E11 and continues throughout development. Different mechanisms contribute to this process including cochlear progenitor proliferation and convergent extension. We previously identified that FGF9 and FGF20 promote cochlear lengthening by regulating auditory sensory epithelial proliferation through FGFR1 and FGFR2 in the periotic mesenchyme. Here, we provide evidence that ETS-domain transcription factors ETV4 and ETV5 are downstream of mesenchymal FGF signaling to control cochlear lengthening. Next generation RNA sequencing identified that *Etv1, Etv4* and *Etv5* mRNAs are decreased in the *Fgf9* and *Fgf20* double mutant periotic mesenchyme. Deleting both *Etv4* and *Etv5* in periotic mesenchyme resulted in shortening of cochlear length but maintaining normal patterning of organ of Corti and density of hair cells and supporting cells. This recapitulates phenotype of mesenchymal-specific *Fgfr1* and *Fgfr2* deleted inner ear. Furthermore, analysis of *Etv1/4/5* triple conditional mutants revealed that ETV1 does not contribute in this process. Our study reveals that ETV4 and ETV5 function downstream of mesenchymal FGF signaling to promote cochlear lengthening.

## 1. Introduction

In the mouse, a ventral outgrowth from the otocyst at approximately E11 marks the developing cochlear duct. Over the following 5 days, the cochlear duct elongates and coil until it reaches one and three-quarters turns (Morsli et al., 1998). Different factors contribute to such elongation including proliferation of a population of progenitor cells located in the floor of the developing cochlear duct. Such population becomes evident early around E11 and is marked with the expression of Sox2. These progenitor cells proliferate for around 48 hours then undergo cell cycle exit, starting in the apex of the cochlear duct around E13 (Doetzlhofer et al., 2006; Matei et al., 2005). Mechanisms regulating the proliferation and cell cycle exit of cells within the progenitor population are under investigation. In addition to progenitor proliferation, convergent extension contributes to the elongation of the sensory organ of the cochlea through unidirectional extension (Wang et al., 2005).

The E26 transformation-specific (ETS) proteins are a group of transcription factors that are encoded by 28 different genes in humans (Oh et al., 2012). These proteins are characterized by the ETS domain which is a highly conserved DNA binding domain. The human ETS proteins are clustered into 12 subgroups based on structural and functional similarity. The PEA3 subgroup includes 3 members: ETV1 (ER81), ETV4 (PEA3) and ETV5 (ERM) which are more than 95% identical in the amino acid sequence within the DNA-binding domain (de Launoit et al., 1997). Members of this subfamily are expressed in multiple organs during development including limb buds, mammary gland and kidney where they are required for normal development (Chotteau-Lelievre et al., 1997; Lu et al., 2009; Mao et al., 2009). Multiple studies show this group of transcription factor function downstream FGF during development (Firnberg and Neubuser, 2002; Herriges et al., 2015; Mao et al., 2009).

FGF signaling plays diverse roles during cochlear development (Ebeid and Huh, 2017). We previously showed that FGF9 and FGF20 are expressed in the epithelium of otic vesicle and signal to the surrounding mesenchymal FGFR1 and FGFR2 to promote cochlear sensory progenitor proliferation and subsequent cochlear growth (Huh et al., 2015). Through both gain- and loss-of-function experiments, we showed that mesenchymal FGF signaling is both necessary and sufficient for cochlea lengthening. To date, the molecules downstream of mesenchymal FGF signaling regulating cochlear length is not known. Here, we identify and validate differentially expressed genes within the periotic mesenchyme of *Fgf9/20* double mutants including three ETV transcription factors; *Etv4, Etv5* and *Etv1*. In addition, deleting *Etv4* and *Etv5* in periotic mesenchyme results in cochlear shortening. These results demonstrate that ETV4 and ETV5 function downstream mesenchymal FGF signaling to regulate cochlear length.

## 2. Materials and methods

### 2.1 Animals

This study was carried out in accordance with the recommendations in the Guide for the Care and Use of Laboratory Animals of the National Institutes of Health. The protocol was approved by the University of Nebraska Medical Center Institutional Animal Care and Use Committee (16-005-02-EP). All efforts were made to minimize animal suffering. *Fgf9*^*-/+*^ (Colvin et al., 2001), *Fgf20*^*−/+*^ (Huh et al., 2012), and *Twist2*^*Cre/+*^ (Sosic et al., 2003) mouse lines were reported previously. *Etv1*^*fl/+*^ (Patel et al., 2003), *Etv4*^*-/+*^ (Laing et al., 2000), and *Etv5*^*fl/+*^ (Zhang et al., 2009) were gifts from Drs. Silvia Arber from University of Basel, John Hassell from McMaster University, and Xin Sun from University of Wisconsin-Madison, respectively. *Fgf9*^*-/+*^;*Fgf20*^*-/+*^ males were mated with *Fgf9*^*-/-*^;*Fgf20*^*-/-*^ females to generate control (*Fgf9*^*-/+*^;*Fgf20*^*-/+*^) and *Fgf9*^*-/-*^ ;*Fgf20*^*-/-*^ embryos. *Etv4*^*-/-*^;*Etv5*^*fl/fl*^;*Twist2*^*Cre/+*^ embryos were generated by crossing *Etv4*^*-/+*^;*Etv5*^*fl/+*^;*Twist2*^*Cre/+*^ males to *Etv4*^*-/-*^;*Etv5*^*fl/fl*^ females. *Etv1*^*fl/fl*^;*Etv4*^*-/-*^;*Etv5*^*fl/fl*^;*Twist2*^*Cre/+*^ were generated by crossing *Etv1*^*fl/+*^;*Etv4*^*-/+*^;*Etv5*^*fl/+*^;*Twist2*^*Cre/+*^ males to *Etv1*^*fl/fl*^;*Etv4*^*-/-*^;*Etv5*^*fl/fl*^ females. Mice were maintained on a 129×1/SvJ;C57B6/J mixed background. Animals for timed mating were put together in the evening, and each morning were tested for the presence of the vaginal plug then they were considered as embryonic day 0.5 (E0.5).

### 2.2 Sample collection and tissue preparation

For laser capture microdissection and RNA sequencing, E11.5 and E12.5 embryos carrying *Fgf9*^*-/-*^;*Fgf20*^*-/-*^ along with control (*Fgf9*^*-/+*^;*Fgf20*^*-/+*^) were collected in DEPC-treated cold PBS. The whole head was immediately embedded in OCT, then frozen in liquid nitrogen and stored at - 80°C until further processing.

### 2.3 Laser capture microdissection

OCT-embedded samples from *Fgf9*^*-/-*^;*Fgf20*^*-/-*^ and littermate controls were serially sectioned horizontally (10µm thick) at -20°C using a cryostat (Leica CM1950). The sections were mounted on slides covered with a Polyethylene Naftelato (PEN) membrane (Arcturus, LCM0522). Slides were allowed to dehydrate inside the cryostat chamber for 10 minutes. Slides were immediately stained with a rapid protocol for eosin on ice. Briefly, tissue sections were fixed in 70% ethanol for 1 minute, hydrated in DEPC-treated water twice for 30 seconds, dehydrated in 95% ethanol for 30 seconds and then stained with an eosin Y solution in 95% ethanol for 30 seconds. Sections were then washed twice in 95% ethanol then twice in 100% ethanol. Finally, sections were dried at room temperature for 1 minute and immediately processed with the microdissection system. Laser capture microdissection (LCM) was performed using a PALM MicroBeam system (Zeiss) as demonstrated in the manual. Specifically, the cochlear duct was visualized under bright-field with a 40x objective. The mesenchyme adjacent to the sensory side of the cochlear duct was selected using the PALM RoboSoftware then cut and catapulted by laser pulses into a microtube cap filled with 30µl of RNA extraction buffer. The capture success was evaluated by checking the slide before and after the microdissection process and observing the captured tissue in the microtube cap.

### 2.4 RNA extraction and quality assessment

Arcturus PicoPure RNA isolation kit (applied biosystems, 12204-01) was used for all RNA extractions as per the manufacturer protocol. Briefly, microdissected tissues in RNA extraction buffer were incubated at 42°C for 30 minutes then stored at -80°C until completing all sample collection. Three littermates per genotype per developmental time point were pooled. During RNA extraction, columns were treated with RNase-free DNase I set (Qiagen, 79254) to remove genomic DNA. The yield and integrity of total RNA from microdissected samples were measured using a 2100 Bioanalyzer (Agilent Technologies). A RIN value was obtained for each sample on a scale of 1–10 and the RNA quality is considered good if RIN ≥ 5 (Kerman et al., 2006).

### 2.5 RNA sequencing and transcriptome analysis

For LCM samples, next generation RNA sequencing was performed at University of Nebraska Medical Center Sequencing core. Briefly, synthesis of cDNA from 1–3ng of total RNA was performed with the Clontech SMARTer™ Ultra Low RNA Kit (Clontech Laboratories Inc, Mountain View, CA). Sequencing libraries were prepared using the Illumina Paired End Sample Prep Kit (Illumina Inc, San Diego, CA) according to Illumina’s Ultra Low Input mRNA-Seq protocol. Each cDNA library was sequenced to generate 75 base pair reads, on the Illumina NextSeq500 (Illumina). The sequence reads were aligned to the mouse reference genome sequence (USCS mm10) using STAR aligner (Dobin et al., 2012). Alignments were assembled and annotated using Cufflinks (Trapnell et al., 2010). DESeq2 (Anders and Huber, 2010) was used to detect differentially expressed gene transcripts.

### 2.6 Pathway analysis

The database DAVID (the database of annotation, visualization and integrated discovery (Huang da et al., 2009a, b) was used for pathway analysis. The differentially expressed gene transcripts (q <0.05) identified from the RNA-Seq data were input into DAVID, which identified enriched biological pathways.

### 2.7 RNA *in situ* hybridization

E11.5 and E12.5 embryos were collected in DEPC-treated cold PBS, fixed with 4% paraformaldehyde overnight at 4°C then washed with PBS three times. RNA *in situ* hybridization analysis was performed on 7µm paraffin sections following a standard procedure with digoxigenin-labeled antisense riboprobes (Moorman et al., 2001). Probes used were as follows: *Etv4* (NM_007424), *Etv5* (NM_001358428.1), *Etv1* (NM_007960.5), *Fam19a4* (NM_177233.5), *Tgfbi* (NM_009369.5), and *Ebf1* (NM_001290709.1).

### 2.8 Quantitative real time PCR

E11.5 and E12.5 embryos were collected, whole cochlear part of the inner ear was dissected out in DEPC-treated cold PBS, then placed in RNA extraction buffer. RNA was extracted using Arcturus PicoPure RNA isolation kit (applied biosystems, 12204-01) according to the manufacturer’s protocol. During RNA extraction, columns were treated with RNase-free DNase I set (Qiagen, 79254) to remove genomic DNA. Reverse transcription of total RNA was performed using GoScript™ Reverse Transcription System (Promega) according to the manufacturer’s protocol. Quantitative real time PCR was performed using both PowerUp™ SYBR™ Green (Applied Biosytems, A25742) and TaqMan Fast Advanced Master Mix (Applied Biosytems, 4444556) according to the manufacturer’s protocol. Data analysis was performed using the comparative CT method, and data were normalized to detection of GAPDH and 18S ribosomal RNA. Primers and probe sets are included in supplemental table S2.

### 2.9 Immunostaining

For whole mount immunostaining, P0 pups carrying individual or combined *Etv1/4/5* mutation along with littermate controls were collected, inner ears were dissected in cold PBS, fixed with 4% paraformaldehyde overnight at 4°C then washed with PBS three times and blocked with PBS containing 0.1% Tween 20 and 0.5% donkey serum. Samples were incubated in primary antibody overnight at 4°C. Samples were then washed with PBS and incubated with a secondary antibody for 2hrs at room temperature then washed, placed on a glass microscope slide in 95% glycerol, coverslipped, and photographed using a Zeiss LSM 700 confocal microscope. Primary antibodies/stains used: Phallodin (R&D Systems, 1:40), Prox1 (Covance, 1:250), Myo6 (Proteus, 1:200).

### 2.10 Cochlear length quantification and cell counting

Cochlear length was measured using FIJI software from the tip of the apex to the base. Myosin 6 and Prox1 immunostaining were used as markers for hair cells and supporting cells respectively. To measure the density, at least 400µm regions of the base, middle, and apex of the cochleae were counted and normalized to 100µm. Cell counting was performed using FIJI software.

### 2.11 Statistics

For each experiment, the numbers of samples (n) is indicated. The p value for difference between samples was calculated using either multiple-testing adjusted p-value (for differential expression using DESeq2 (Anders and Huber, 2010)) or a two-tailed Student’s t-test (for the qRT-PCR and length quantification), and p < 0.05 was considered as significant.

## 3. Results

### 3.1 Gene Expression Profiling reveals potential FGF-regulated genes of periotic mesenchyme

We previously identified that mesenchymal FGF signaling is necessary and sufficient for auditory sensory progenitor proliferation and subsequent cochlear lengthening (Huh et al., 2015). To identify genes regulated by FGF signaling in periotic mesenchyme, we isolated E11.5 and E12.5 mesenchymal tissues adjacent to the developing cochlear duct from *Fgf9*^*-/-*^;*Fgf20*^*-/-*^ and control (*Fgf9*^*-/+*^;*Fgf20*^*-/+*^) embryos using laser-capture microscopy. We then performed next-generation RNA sequencing. Analysis of normalized sequence reads in control samples indicated enrichment of periotic mesenchymal genes such as *Pou3f4* and *Tbx18*, while epithelial genes like *Sox2* and *Fgf20* exhibited very low sequence reads (supplemental table S1). Such results provide evidence of the purity of the collected cell population using laser capture microdissection and thereby the reliability and reproducibility of our data.

We next analyzed differentially expressed genes at E11.5 comparing *Fgf9*^*-/-*^;*Fgf20*^*-/-*^ to control embryos. In *Fgf9*^*-/-*^;*Fgf20*^*-/-*^ mesenchyme, 48 genes showed upregulation and 6 genes showed downregulation compared to control (Fig. 1A) (Log2 fold change >1, adjusted p-value<0.05). At E12.5, 68 genes showed upregulation and 55 genes showed downregulation (Fig. 1B) (Log2 fold change >1, adjusted p-value<0.05). The number of differentially expressed genes (especially the downregulated genes) increased at E12.5 compared to E11.5 indicating subsequent transcriptome changes occurring in a 24-hour interval. The top downregulated genes at both time points included the ETS-domain transcription factors (*Etv1, Etv4, Etv5*) as well as other transcription factors such as *Ebf1*. Interestingly, chemokine-like secreted protein *Fam19a4* was the most downregulated gene at both time points with more than 4 folds. In addition, *Itga8* gene that encodes the alpha 8 subunit of the heterodimeric integrin alpha8beta1 protein showed downregulation more than 4 folds at E12.5. The top differentially expressed genes at both time points are shown in table 1.

**Table 1.**
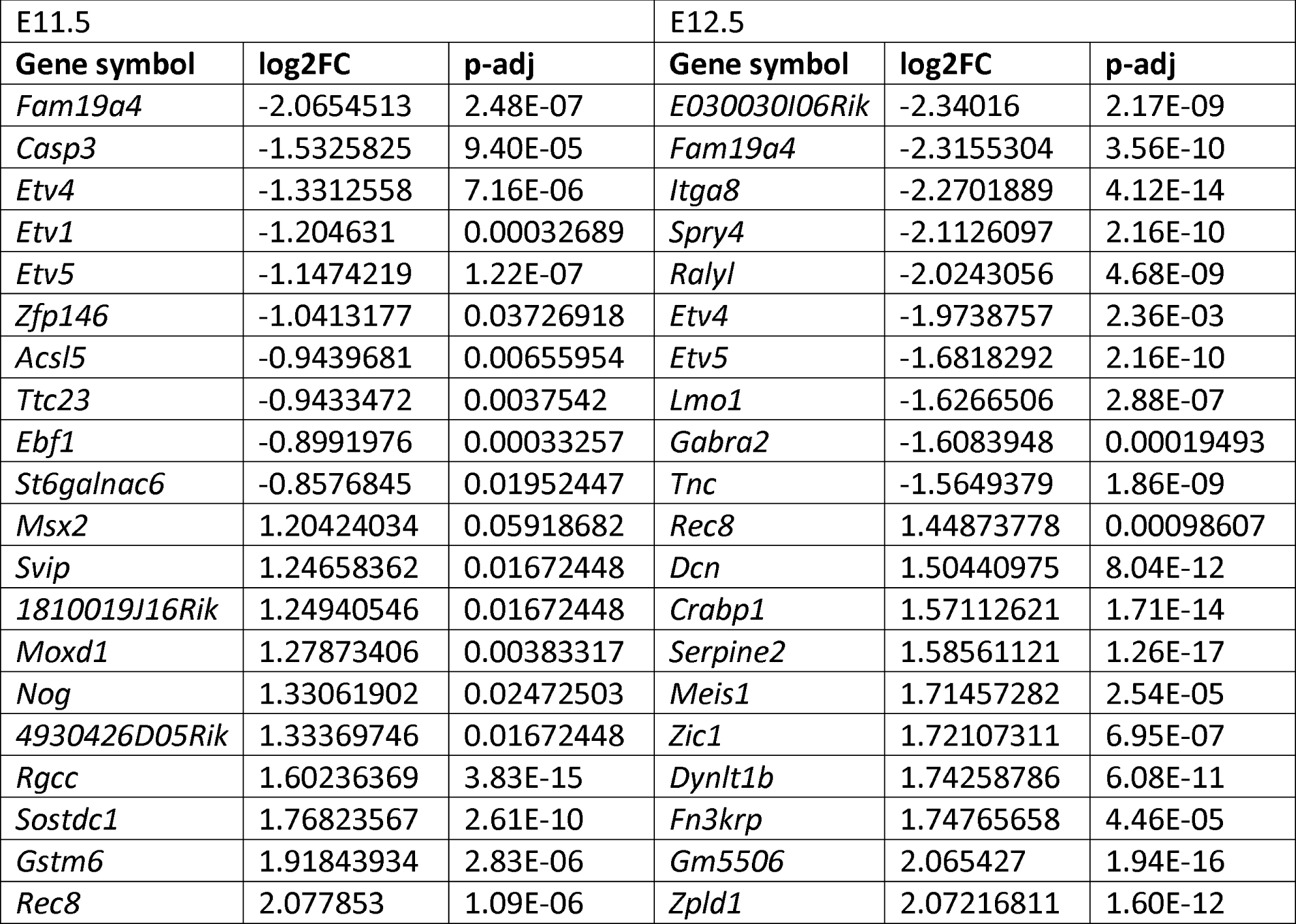
Top differentially expressed genes at E11.5 and E12.5 in *Fgf9*^*-/-*^;*Fgf20*^*-/-*^ versus control. Change is gene expression is shown in log2 fold change (log2FC) and statistical significance is demonstrated as adjusted p-value (P-adj.).

**Fig 1.**
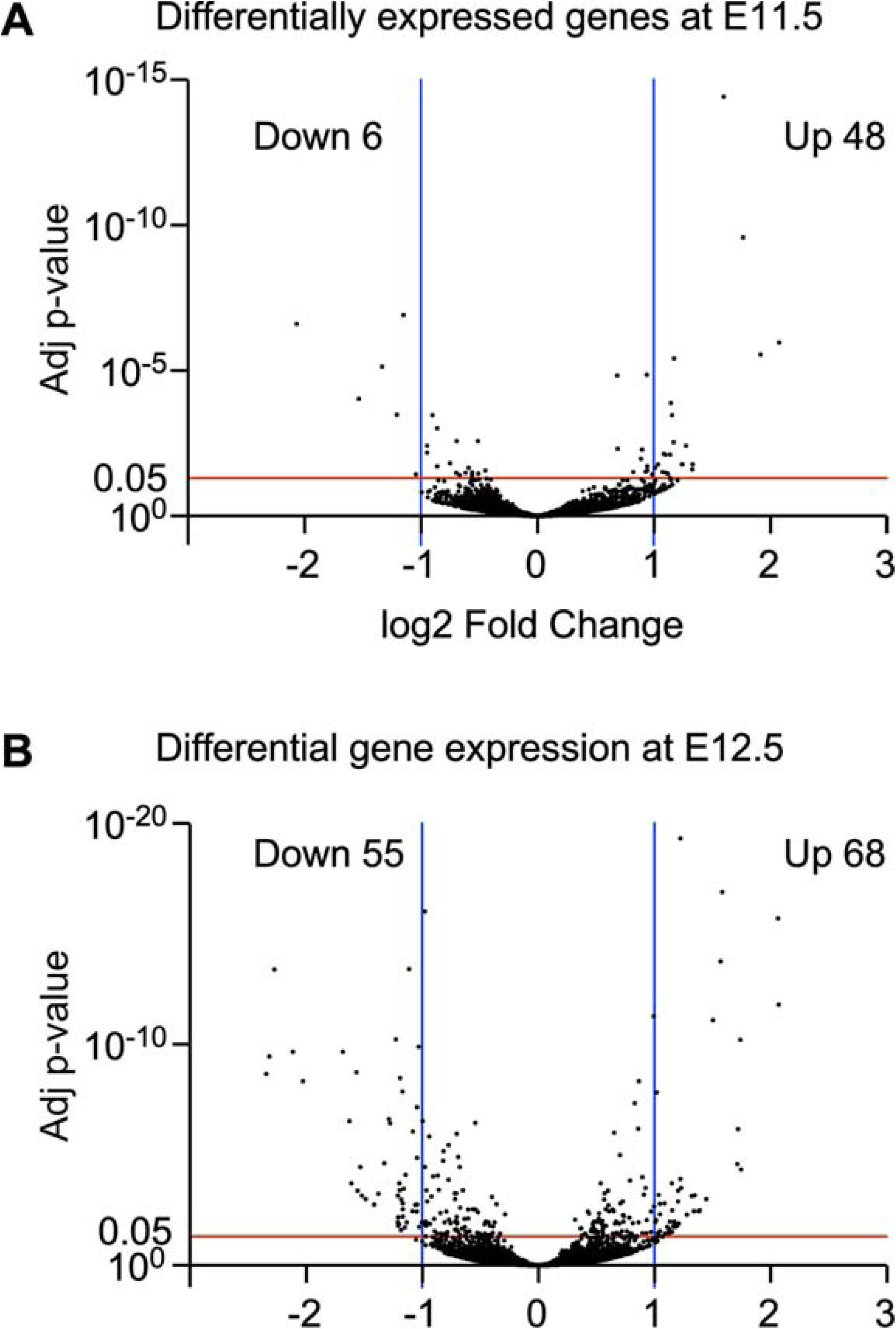
Differentially expressed genes in *Fgf9*^*-/-*^;*Fgf20*^*-/-*^ mesenchyme versus control. Volcano plot showing up- and down-regulated genes at E11.5 (A) and E12.5 (B). Red line indicates the threshold of adjusted p-value (0.05) while the blue lines show the cutoff of log2 fold change (>1 and <-1). Each dot on the plot represents a single gene.

### 3.2 Functional annotation of differentially expressed genes shows enriched pathways/functions

Next, we studied how differentially expressed genes in *Fgf9*^*-/-*^;*Fgf20*^*-/-*^ mesenchyme relates to the development of cochlear progenitors. To understand the functional categories enriched in differentially expressed genes, we utilized the Database for Annotation, Visualization and Integrated Discovery (DAVID) for functional annotation of differentially expressed gene (Huang da et al., 2009a, b) Gene ontological analyses of downregulated genes at both time points showed enrichment in transcription factors, specifically PEA3 group of ETS transcription factors (*Etv1, Etv4 and Etv5*). Additionally, at E11.5, *Ebf1* and *Tfdp1* transcription factors as well as cell cycle regulators (*Ccnd2, Tfdp1*) were downregulated while at E12.5, negative regulators of tyrosine kinase receptor signaling (Spry1 and Spry4) and multiple developmental proteins were downregulated (Table 2).

**Table 2.**
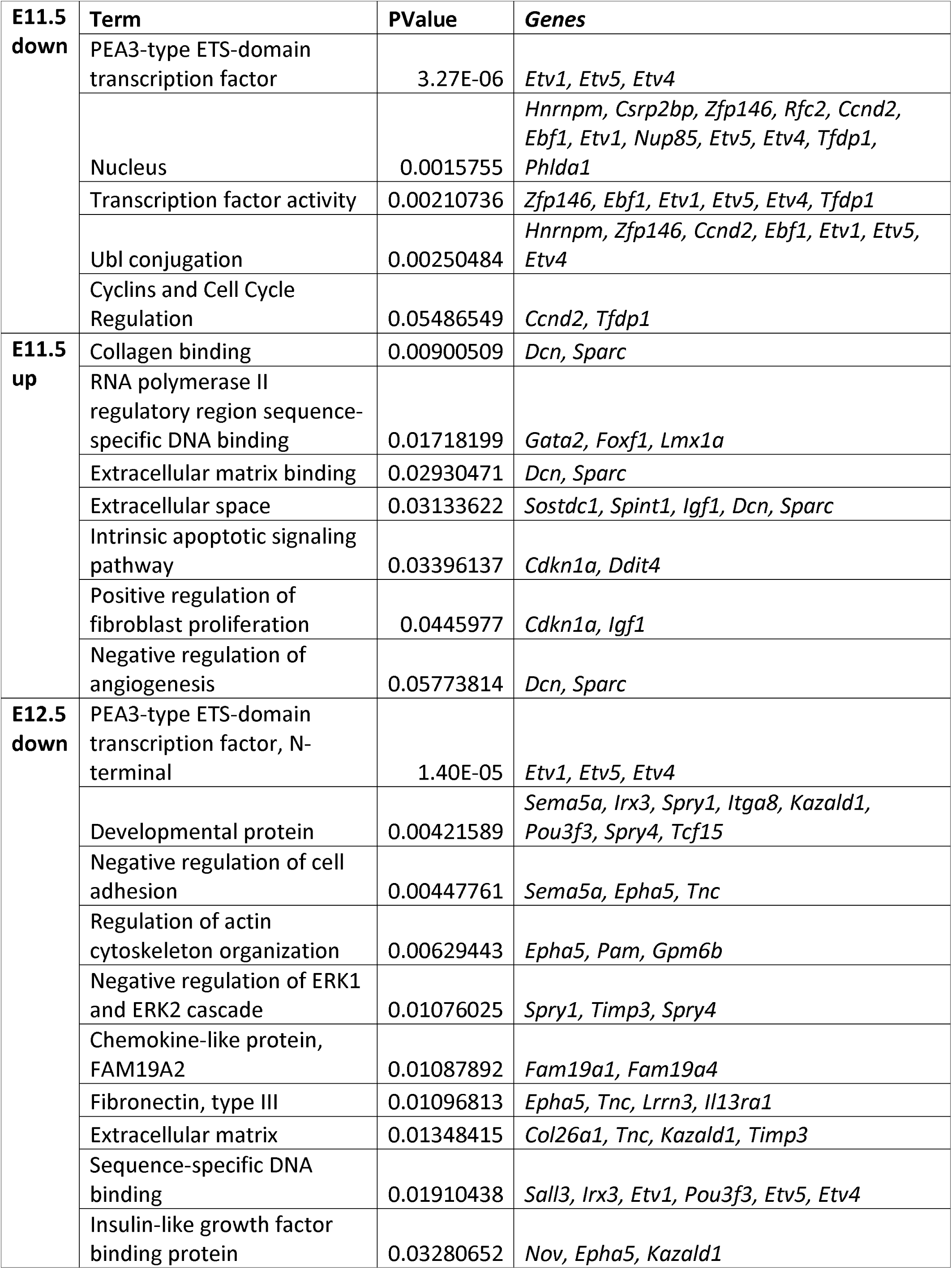
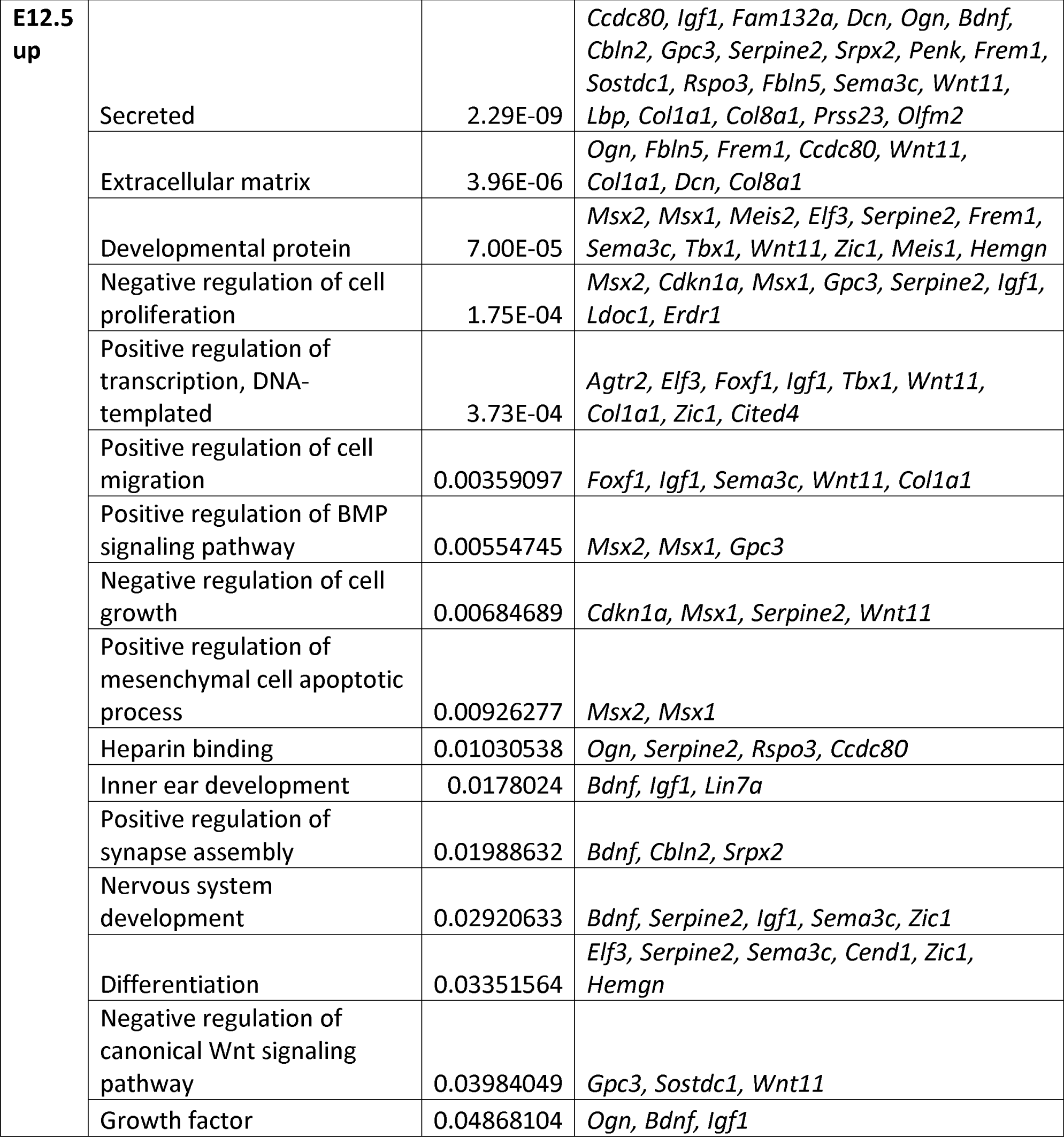
Functional categories of differentially expressed genes (either upregulated (up) or downregulated (down)) at E11.5 and E12.5 in *Fgf9*^*-/-*^;*Fgf20*^*-/-*^ versus control mesenchyme.

As for upregulated gene categories at E11.5, genes encoding collagen binding proteins (*Sparc and Dcn*), as well as other secreted proteins (*Sostdc1, Spint1, and Igf1*) were observed. At E12.5, multiple genes encoding secreted proteins as well as negative regulators of cell proliferation were upregulated (Table 2).

### 3.3 Differentially expressed genes validated through quantitative RT-PCR and *in situ* hybridization

To validate differential gene expression detected by RNA sequencing, quantitative RT-PCR was performed on dissected E11.5 and E12.5 cochlea from *Fgf9*^*-/-*^;*Fgf20*^*-/-*^ and littermate controls. At E11.5, downregulation was verified for *Etv4, Fam19a4, Tgfbi* and *Spry4* while upregulation was verified for *Dcn, Sparc* and *Nog* (n = 3, Student’s t-test p<0.05). *Etv1* and *Etv5* showed downward trends consistent with the RNA sequencing data (Fig. 2A). At E12.5, downregulation was validated for *Etv4, Fam19a4, Tgfbi, Kitl, Tbx18, Pou3f3, Itga8*, and *Enpp2* (n = 3, Student’s t-test p<0.05) (Fig. 2B).

**Fig 2.**
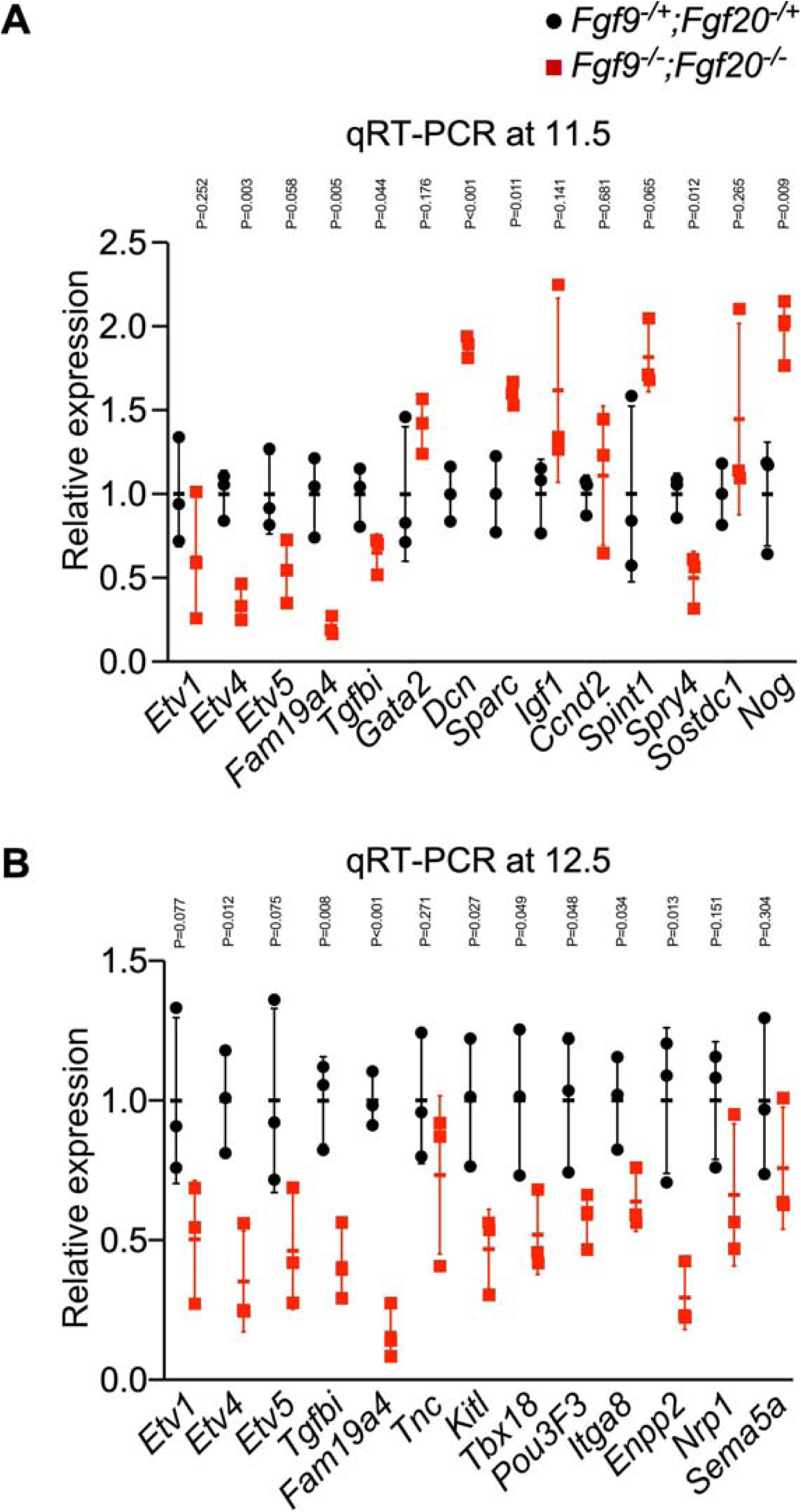
Validating differentially expressed genes in *Fgf9*^*-/-*^;*Fgf20*^*-/-*^ versus control. Linear fold change in gene expression in *Fgf9*^*-/-*^;*Fgf20*^*-/-*^ (red) compared to control (black) at E11.5 (upper panel) and E12.5 (lower panel) (n = 3, Student’s t-test p-value per gene is shown).

Also, *in situ* hybridization was performed probing a subset of differentially expressed genes. Etv4 and Etv5 exhibited a similar expression domain at E11.5 and E12.5 where the signal was detected in the mesenchyme surrounding the cochlear duct with a higher signal on the non-sensory side (Fig. 3). The sensory epithelium showed signals for both genes with slightly different domains. *Etv5* was also detected in the non-sensory epithelium at E12.5 (Fig. 3). As for *Etv1*, its expression was weak compared to *Etv4* and *Etv5*. The signal was only detected in the mesenchyme on the non-sensory side of the cochlea. *Ebf1* and *Tgfbi* were detected in the mesenchyme surrounding the cochlear duct. *Fam19a4* was expressed in the mesenchyme on the non-sensory side of the growing cochlea. All genes analyzed by *in situ* hybridization was downregulated in *Fgf9*^*-/-*^;*Fgf20*^*-/-*^ compared to controls.

**Fig. 3.**
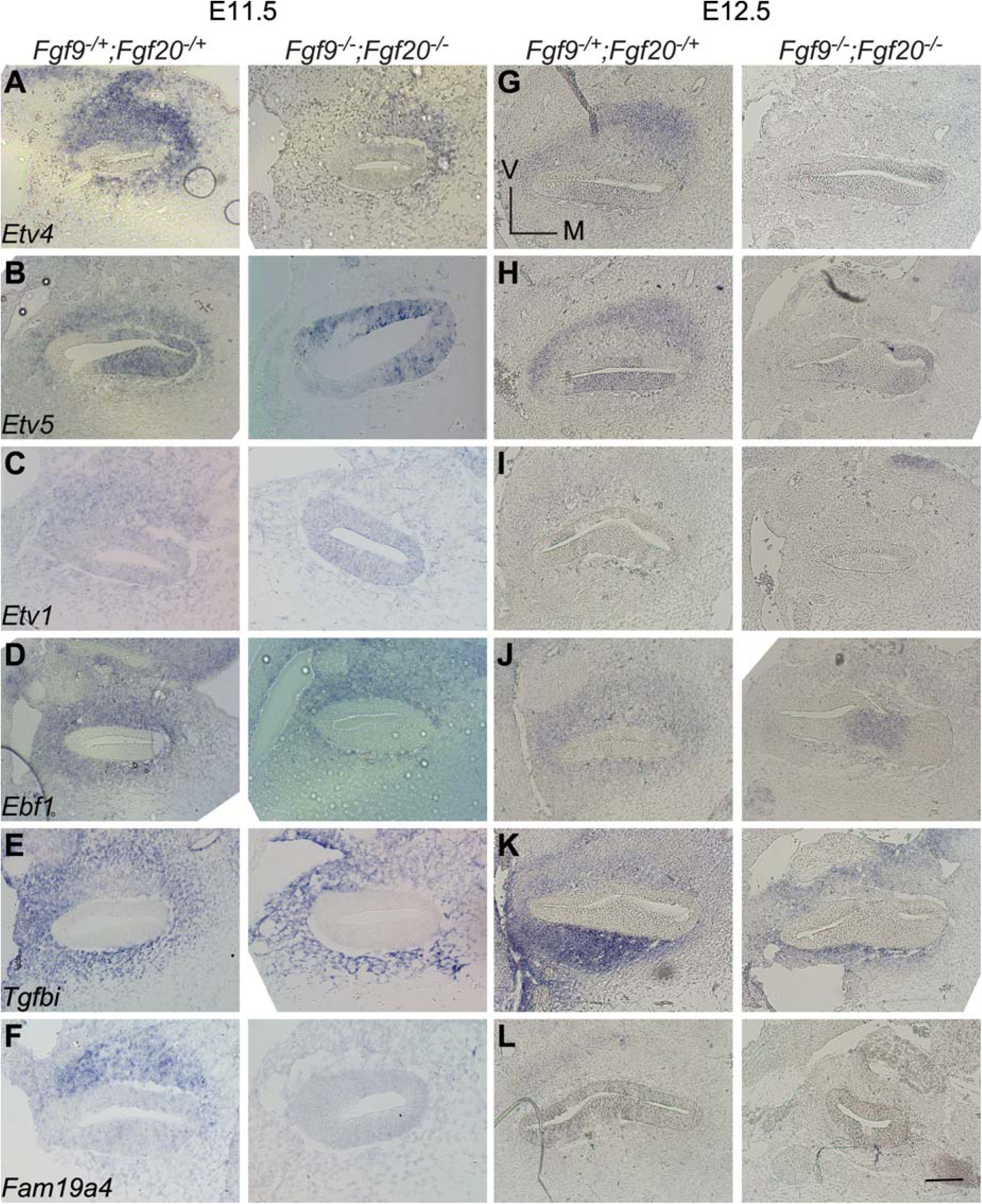
In-situ hybridization of selected candidate genes downstream FGF signaling in periotic mesenchyme at E11.5 and E12.5 in *Fgf9*^*-/-*^;*Fgf20*^*-/-*^ versus control. Scale bar=50μm.

### 3.4 *Etv4* and *Etv5* are required for cochlear lengthening but not epithelial patterning during development

Since *Etv4* and *Etv5* showed downregulation at both time points, and both factors function as downstream effectors of FGF signaling in other organ development (Ornitz and Itoh, 2015), we examined their functions during cochlear development. We used a mesenchymal driver (*Twist2*^*Cre*^) (Sosic et al., 2003) and *Etv5* Floxed mice (*Etv5*^*fl/+*^) (Zhang et al., 2009) to delete *Etv5* in periotic mesenchyme with conventional deletion of Etv4. Examination of P0 cochlea showed that length of cochleae from *Etv4*^*-/-*^;*Etv5*^*fl/+*^;*Twist2*^*Cre/+*^ and *Etv4*^*+/-*^;*Etv5*^*fl/fl*^;*Twist2*^*Cre/+*^ were comparable to controls (Fig. 4). Length of cochleae from *Etv4*^*-/-*^;*Etv5*^*fl/fl*^;*Twist2*^*Cre/+*^ was decreased to 86% compared to controls (Student t-test, n at least 6, p=0.00014) (Fig. 4). These data indicate that both *Etv4* and *Etv5* are required for cochlear lengthening. Next, we examined the patterning and cell density of the organ of Corti in each model. Staining for hair cell stereocilia bundles and supporting cells showed that hair cells and supporting cells align well in all genotypes (Fig. 5). In addition, the densities of hair cells and supporting cells were comparable in all genotypes (Fig. 5). These data indicate that mesenchymal *Etv4* and *Etv5* are dispensable for sensory epithelial patterning and differentiation.

**Fig 4.**
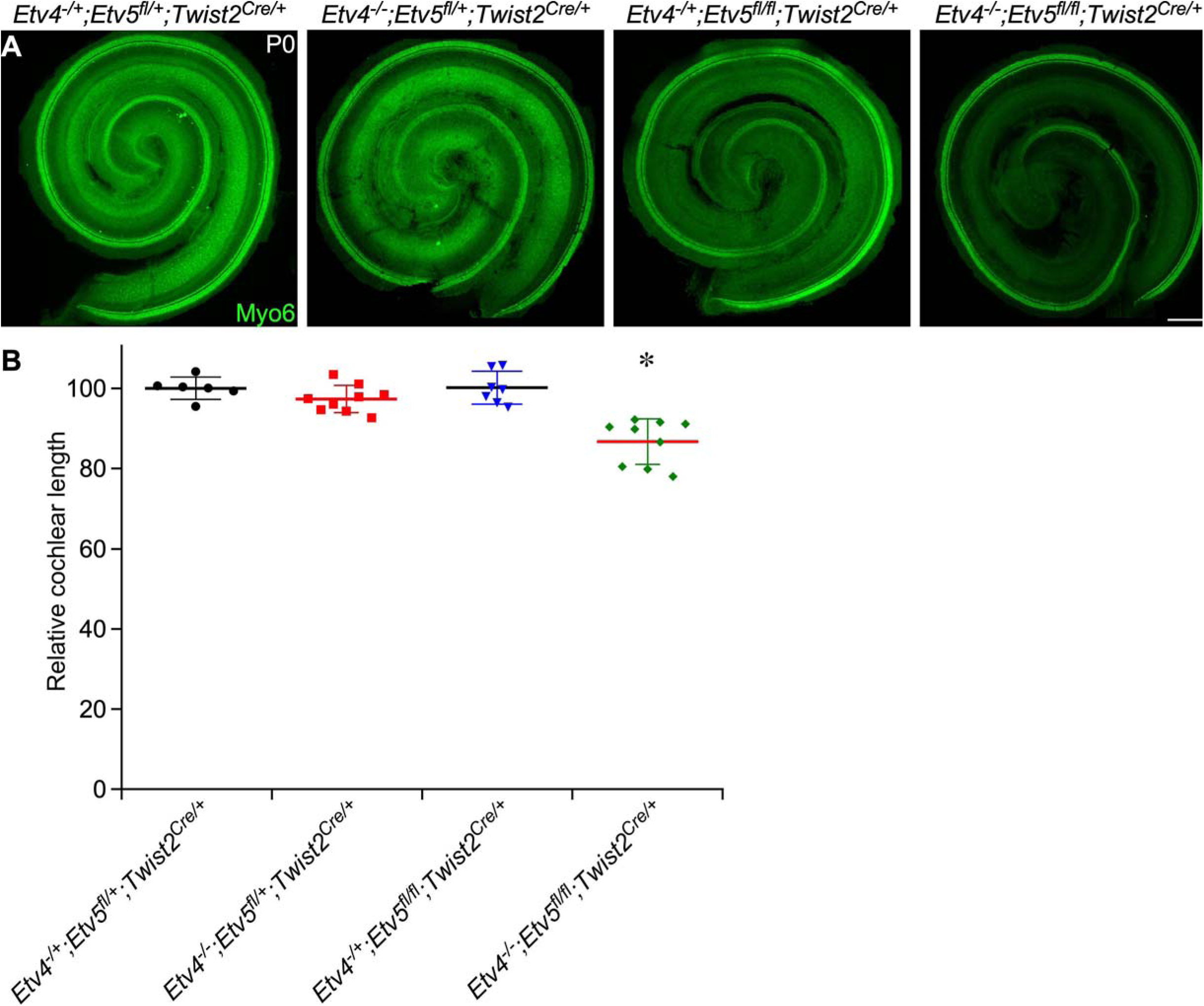
Cochlear phenotype in Etv4/5 compound mutants. (Upper panel) Whole mount cochlea from P0 Etv4 KO (*Etv4*^*-/-*^; *Etv5*^*fl/+*^;*Twist2*^*Cre/+*^), Etv5 cKO (*Etv4*^*-/+*^;*Etv5*^*fl/fl*^;*Twist2*^*Cre/+*^), Etv4/5 compound mutant (*Etv4*^*-/-*^;*Etv5*^*fl/fl*^;*Twist2*^*Cre/+*^) and littermate control (*Etv4*^*-/+*^;*Etv5*^*fl/+*^;*Twist2*^*Cre/+*^) showing whole cochlea stained with Myosin 6 antibody (green). (Lower panel) Quantification of cochlear duct length at P0 from each mouse model relative to control. Scale bar=200μm.

**Fig 5.**
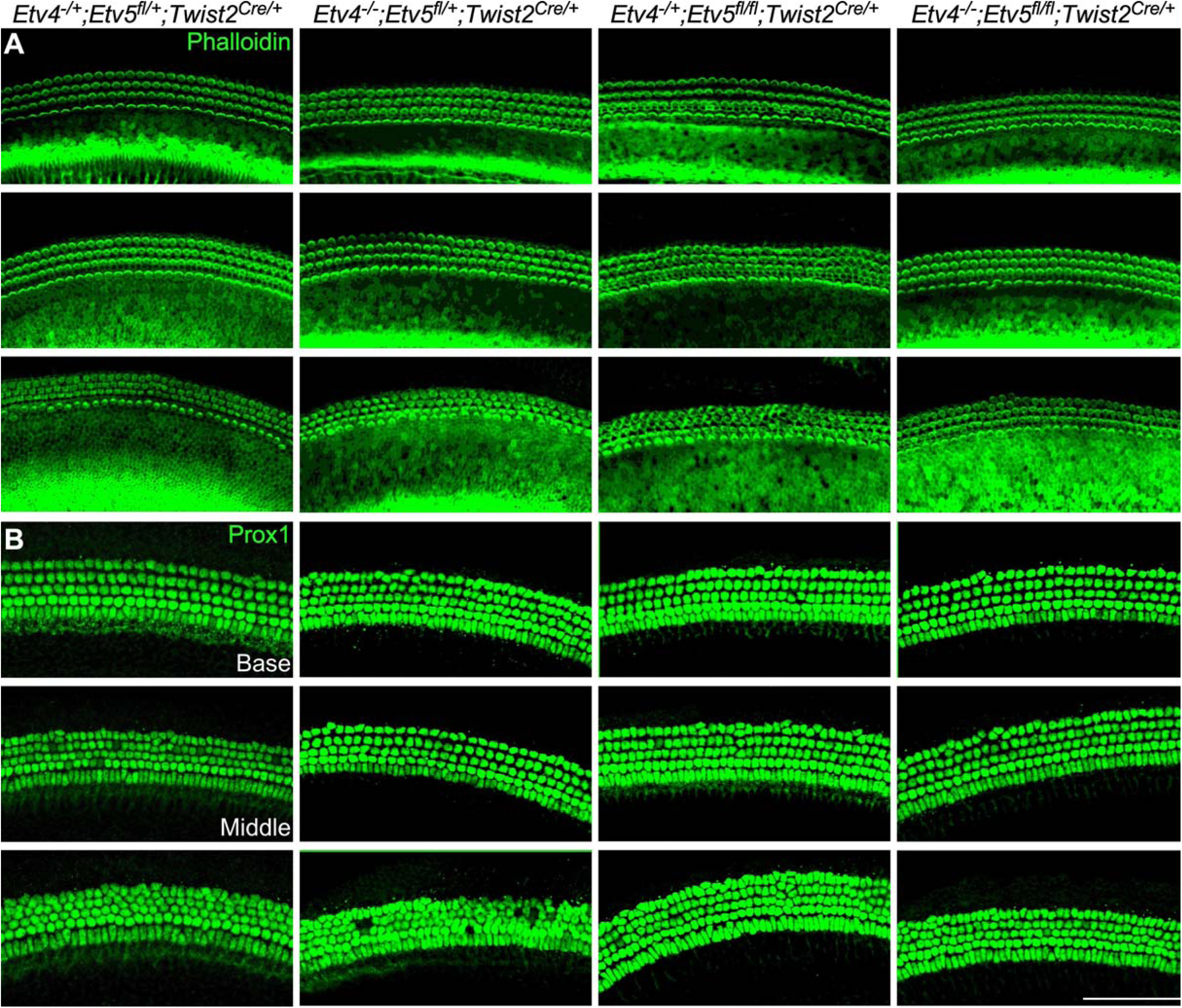
Hair cells and supporting cells in Etv4/5 compound mutants. Whole mount immunostaining of P0 cochlea from Etv4 KO (*Etv4*^*-/-*^; *Etv5*^*fl/+*^;*Twist2*^*Cre/+*^), Etv5 cKO (*Etv4*^*-/+*^;*Etv5*^*fl/fl*^;*Twist2*^*Cre/+*^), *Etv4/5* compound mutant (*Etv4*^*-/-*^;*Etv5*^*fl/fl*^;*Twist2*^*Cre/+*^) and littermate control (*Etv4*^*-/+*^;*Etv5*^*fl/+*^;*Twist2*^*Cre/+*^) showing representative regions from each cochlear turn stained with F-actin to show Actin-enriched hair cell stereocilia (green) in the upper row, and Prox1 staining (green) in the lower row as a marker for supporting cells. Scale bar=100μm.

**Fig 6.**
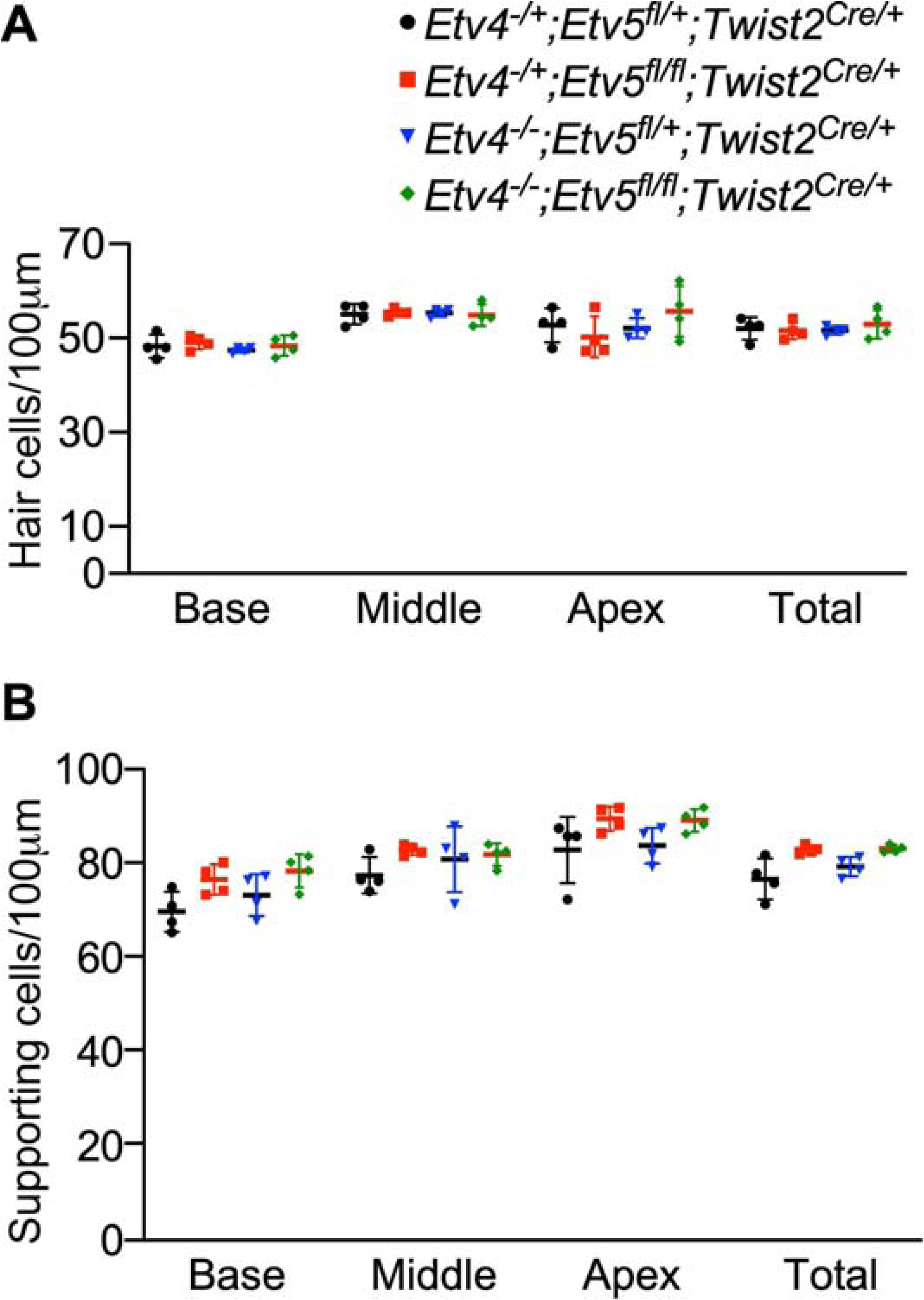
Hair cell and supporting cell density in *Etv4/5* compound mutants Quantification of hair cell and supporting cell density at P0 from Etv4 KO (*Etv4*^*-/-*^; *Etv5*^*fl/+*^;*Twist2*^*Cre/+*^), *Etv5* cKO (*Etv4*^*-/+*^;*Etv5*^*fl/fl*^;*Twist2*^*Cre/+*^), Etv4/5 compound mutant (*Etv4*^*-/-*^ ;*Etv5*^*fl/fl*^;*Twist2*^*Cre/+*^) and littermate control (*Etv4*^*-/+*^;*Etv5*^*fl/+*^;*Twist2*^*Cre/+*^) showing no significant change.

### 3.5 Mesenchymal Etv1 is dispensable for cochlear development

Compared to the shortened cochlear phenotype caused by the loss of mesenchymal FGF signaling (about 40% compared to controls (Huh et al., 2015)), the phenotype of *Etv4*^*-/-*^ ;*Etv5*^*fl/fl*^;*Twist2*^*Cre/+*^ (86% compared to controls) is moderate. *Etv1* is another member of Erm subfamily of ETS-domain transcription factor (de Launoit et al., 1997). Next-generation RNA sequencing showed that *Etv1* expression was also decreased at both E11.5 and E12.5 in *Fgf9*^*-/-*^ ;*Fgf20*^*-/-*^ mesenchyme tissues compared to controls (Tables 1&2). Therefore, we reasoned that Etv1 may function together with *Etv4* and *Etv5* to regulate cochlear length. To investigated whether *Etv1* functions redundantly with *Etv4* and *Etv5* in regulating cochlear length, *Etv1/4/5* triple conditional mutants were generated. Analysis of the length of the cochleae at P0 showed that shortened cochlear length in *Etv1*^*fl/fl*^; *Etv4*^*-/-*^;*Etv5*^*fl/fl*^;*Twist2*^*Cre/+*^ (83% compared to controls) is comparable to either *Etv1*^*fl/+*^; *Etv4*^*-/-*^;*Etv5*^*fl/fl*^;*Twist2*^*Cre/+*^ (84.5% compared to control) or *Etv4*^*-/-*^ ;*Etv5*^*fl/fl*^;*Twist2*^*Cre/+*^ (86% compared to controls) (Student t-test, n at least 6, p=0.223) (Fig.7A & 8A). Epithelial patterning of organ of Corti and densities of both hair cells and supporting cells were comparable in all genotypes (Fig. 7B,C & 8B,C) indicating that mesenchymal Etv1 is dispensable for cochlear development.

**Fig 7.**
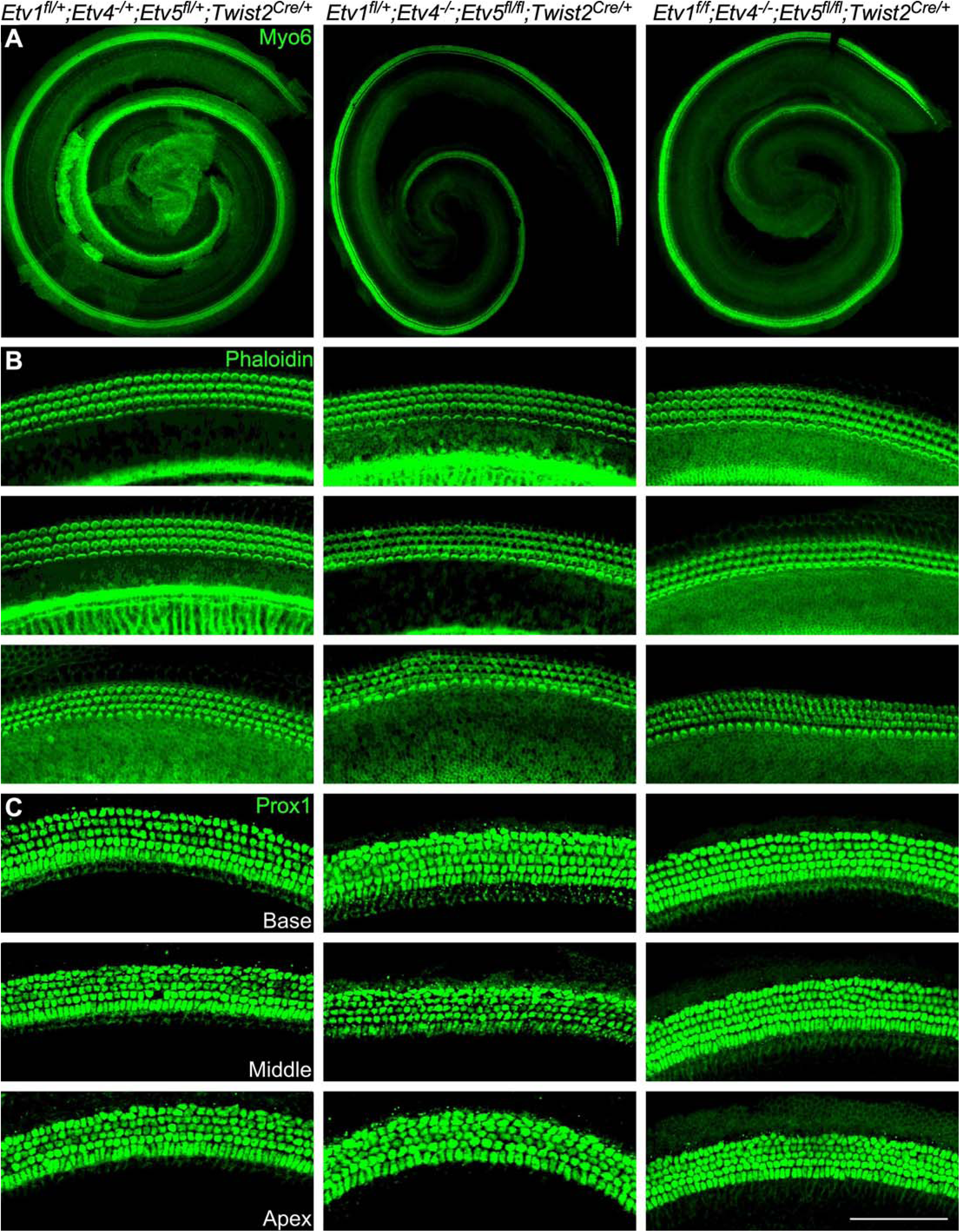
Mesenchymal Etv1 is dispensable for cochlear development Whole mount immunostaining of P0 cochlea from control (*Etv1*^*fl/+*^; *Etv4*^*-/+*^; *Etv5*^*fl/+*^;*Twist2*^*Cre/+*^), *Etv4/5* double mutant (*Etv1*^*fl/+*^;*Etv4*^*-/-*^;*Etv5*^*fl/fl*^;*Twist2*^*Cre/+*^) and *Etv1/4/5* triple mutant (*Etv1*^*fl/fl*^;*Etv4*^*-/-*^;*Etv5*^*fl/fl*^;*Twist2*^*Cre/+*^) showing whole cochlea stained with Myosin 6 antibody (green) (A), and representative regions from each cochlear turn stained with F-actin (green) (B), and Prox1 staining (green) (C) as a marker for supporting cells. Scale bar=100μm.

**Fig 8.**
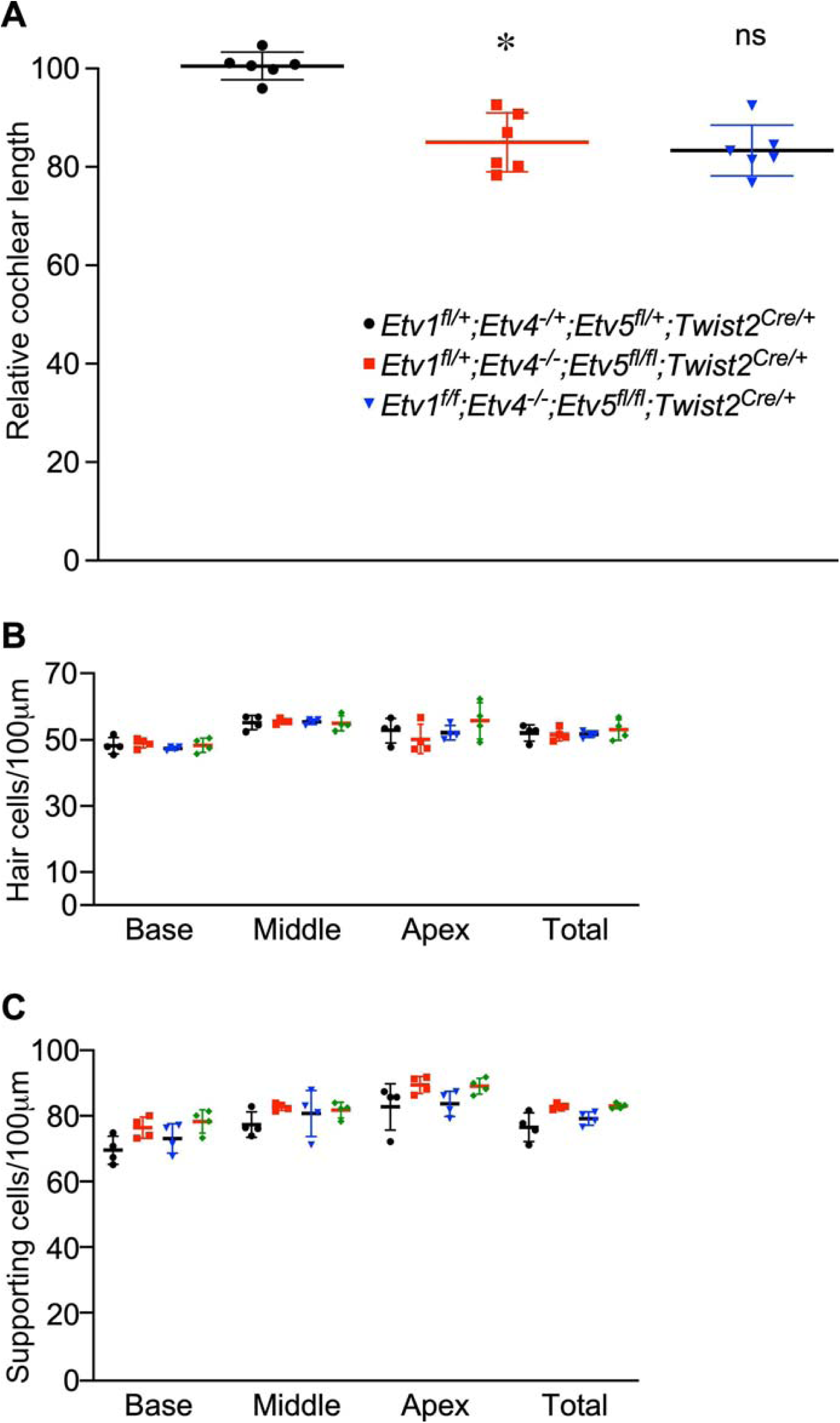
Quantification of cochlear length, hair cell and supporting cell density in *Etv1/4/5* compound mutants (A) Quantification of cochlear duct length at P0 from control (*Etv1*^*fl/+*^;*Etv4*^*-/+*^;*Etv5*^*fl/+*^;*Twist2*^*Cre/+*^), *Etv4/Etv5* double mutant (*Etv1*^*fl/+*^;*Etv4*^*-/-*^;*Etv5*^*fl/fl*^;*Twist2*^*Cre/+*^) and *Etv1/4/5* triple mutant (*Etv1*^*fl/fl*^;*Etv4*^*-/-*^;*Etv5*^*fl/fl*^;*Twist2*^*Cre/*+^) (B) Hair cell density and (C) supporting cell density from each genotype.

## 4. Discussion

### 4.1 FGF signaling regulate the periotic mesenchyme transcriptome

Gene expression analysis of the periotic mesenchyme using RNA sequencing, RNA *in situ* hybridization, and quantitative real time PCR identified multiple targets downstream FGF signaling at E11.5 and E12.5 developmental time points. We looked specifically at 3 sets of genes: transcription factors that function as direct downstream targets of FGF signaling, secreted molecules that can diffuse back to the growing cochlear epithelium to control the developing progenitors, and cell surface molecules that can regulate epithelial-to-mesenchymal interaction.

As for transcription factors, we identified 3 members of the ETS family of transcription factors that belongs to the same PEA3 subfamily; *Etv1, Etv4* and *Etv5* (de Launoit et al., 1997) to be downregulated. In addition, other transcription factors such as *Ebf1* and *Tfdp1* were identified. Such data indicates that FGF signaling at this time of development is regulating mesenchymal gene expression through multiple transcription factors. Another set of downstream targets identified is the Sprouty proteins (*Spry1* and *Spry4*), which are expected as these proteins are known as negative feedback regulators of receptor tyrosine kinase signaling, including FGF signaling (Shim et al., 2005).

Examining genes encoding secreted proteins that were differentially expressed in *Fgf9/20* double mutants revealed *Fam19a4*, a member of the family of chemokine-like proteins, is on the top of downregulated genes at both E11.5 and E12.5 that was also confirmed with real time PCR. Although recent study shows its rule in the activation of macrophages during pathogenic infections (Wang et al., 2015), the role of these proteins during inner ear development is still to be discovered. It is not clear whether FAM19A4 can diffuse back to the developing cochlear epithelium and function to regulate progenitor proliferation. Lack of a mouse model with *FAM19a4* deletion is limiting such investigation. Our data also show other genes encoding secreted proteins to be dysregulated in *Fgf9/20* double mutants such as *Sostdc1*. It is a known antagonist for bone morphogenic protein (BMP) signaling and Wnt signaling pathways (Faraahi et al., 2019) and it can potentially diffuse and regulate epithelial development.

As for the cell surface molecules, we identified *Itga8* gene expression to be downregulated in *Fgf9/20* double mutant mesenchyme at E12.5. As a member of integrin family, this protein functions as a cell adhesion receptor that serves as a link between the extracellular matrix and the cytoskeleton and is capable of bidirectional signaling across the plasma membrane (Hynes, 2002). Follow up studies on the specific rule of such secreted proteins can potentially uncover their contribution to cochlear lengthening.

### 4.2 ETV transcription factors operate downstream FGF signaling to regulate cochlear length

Multiple studies show ETV transcription factor function downstream FGF during development (Firnberg and Neubuser, 2002; Herriges et al., 2015; Mao et al., 2009). Our analysis of *Etv4/5* compound mutant cochleae showed that deletion of *Etv4* or *Etv5* alone maintained normal cochlear length and sensory epithelial patterning. On the other hand, deleting both *Etv4* and *Etv5* yielded a reduced cochlear length. This indicate that *Etv4* and *Etv5* function redundantly during cochlear development. However, the degree of phenotype in *Etv4/5* double mutants (about 86% of the control cochlea) was yet not to the extent of the shortening observed in *Fgf9/20* double mutant (about 40% of the control length) (Huh et al., 2015). This is indicative of other factors that are still expressed in *Etv4/5* double mutants and are capable of compensating the loss of both *Etv4* and *Etv5*. Investigating the role of *Etv1* in cochlear development redundantly with *Etv4/5* didn’t show any significant contribution (Fig. 7). Further experiments investigating other candidate transcription factor such as *Ebf1* contribution in cochlear lengthening are valid future directions.

### 4.3 Mesenchymal ETV transcription factors are dispensable for sensory epithelium patterning and differentiation

Examining hair cells and supporting cells in *Etv4/5* and *Etv1/4/5* compound mutants showed normal patterning and density across the length of the cochlear duct in all genotypes analyzed. Although the density is not changed, the total numbers of hair cells and supporting cells are reduced due to shorter cochlea in *Etv4/5* double mutants and *Etv1/4/5* triple mutants. This is a clear recapitulate of phenotype in mesenchymal *Fgfr1/2* deletion. Whether shortened cochlear impacts hearing function is interesting subject for further investigation.

### 4.4 Future Perspectives

Although our study shows 2 ETV transcription factors regulating the cochlear length, the mechanism of such regulation is still unclear. Gene expression analysis of the developing cochlear sensory epithelium in mouse models with mesenchymal ETV deletion might help identify target genes downstream ETV4 and ETV5 transcription factors. Additionally, chromatin immunoprecipitation and sequencing (ChIP-seq) assay can be utilized to identify genome-wide DNA binding sites for ETV transcription factors during cochlear development. Loss-of-function experiments utilizing either *in vivo* mouse models if available or *ex vivo* explants with gene manipulation will be valid future direction to confirm ETV target contribution in cochlear lengthening.

A challenge facing future experiments is that FGF signaling and subsequently ETVs are active in both cochlear epithelium and surrounding mesenchyme, and possibly playing different roles in each domain at different developmental stages. Careful experimental planning and data interpretation that take into consideration these different expression domains and different roles of FGF signaling during cochlear development is required to achieve better understanding of the underlying molecular mechanisms.

## Acknowledgements

We thank Drs. Silvia Arber from University of Basel, John Hassell from McMaster University, and Xin Sun from University of Wisconsin-Madison for sharing their mouse models, the University of Nebraska Medical Center Sequencing core facility for help in transcriptome analysis, Ligyeom Ha and Brennan Roche for mouse handling.

## Funding

This work was supported by the NIH DC012825, GM110768, and DHHS 44388-Y3 (to S.-H.H.); and the American Hearing Research Foundation (to M.E.).

## Notes

### Competing Interest Statement

The authors have declared no competing interest.

